# Elevational patterns in two groups of micromoths (Lepidoptera: Pterophoridae, Alucitidae) in tropical forests of Mount Cameroon

**DOI:** 10.1101/2025.08.23.671935

**Authors:** Fernando P. Gaona, Sylvain Delabye, Vincent Maicher, Peter Ustjuzhanin, Vasily N. Kovtunovich, Štěpán Janeček, Mercy Murkwe, Pavel Potocký, Szabolcs Sáfián, Robert Tropek

**Affiliations:** Department of Ecology, Faculty of Science, Charles University, Prague, Czechia; Institute of Entomology, Biology Centre, Czech Academy of Sciences, České Budějovice, Czechia; The Nature Conservancy Gabon, Impasse Edowangani, Libreville, Gabon; Altai State University, Barnaul, Russia; Biological Institute, Tomsk State University, Tomsk, Russia; University of the Free State (UFS), Bloemfontein, Free State, South Africa; Department of Biology, Higher Teachers Training College, University of Bamenda, Bambili, Cameroon; Hungarian Natural History Museum, Department of Zoology, Budapest, Hungary

**Keywords:** Afrotropics, altitudinal patterns, Bergmann’s rule, biodiversity, elevational gradients, Microlepidoptera, moths, Rapoport’s rule

## Abstract

Tropical elevational gradients offer unique insights into ecological processes shaping biodiversity, although micromoths remain severely understudied, especially in the Afrotropics. We analysed species richness, community composition, and functional traits (elevational range size and wingspan) of two micromoth families, many-plumed moths (Alucitidae) and plume moths (Pterophoridae), along a complete forest gradient on Mount Cameroon, an Afrotropical biodiversity hotspot. Alucitidae exhibited a distinct mid-elevation diversity peak, mirroring common patterns in tropical butterflies and moths, whereas Pterophoridae showed an uncommon upslope increase in species richness. Both families demonstrated clear elevational turnover in community composition, indicative of strong environmental filtering. Additionally, community-weighted elevational range size and wingspan increased consistently with elevation, supporting Rapoport’s rule and Bergmann’s cline, respectively. These patterns likely reflect interactions among climatic factors and environmental complexity, though the underlying mechanisms remain unresolved. Our findings reveal the unique communities of both mid- and high-elevation forests, as well as the distinctiveness of species-poor lowland assemblages. This elevational differentiation underscores the need for conservation across the full gradient and the vulnerability of endemic highland taxa to climate-driven range contractions. Comprehensive research on neglected tropical insect groups is urgently needed to better anticipate biodiversity responses to environmental change.

## Introduction

Elevational gradients offer natural laboratories for understanding the ecological and evolutionary drivers of biodiversity patterns, as they compress climatic and environmental variation into short geographic distances (McCain & Grytnes, 2010; Rahbek et al., 2019). In tropical regions, where the majority of global biodiversity is concentrated (e.g. Liang et al., 2022), such gradients are particularly valuable for studying how climate, energy availability, and spatial constraints interact to shape species distributions (Rahbek, 1995; McCain & Grytnes, 2010; Peters et al., 2016; Dolson & Kharouba, 2024). Mountains in these regions not only harbour high species richness but also exhibit exceptional rates of turnover and endemism across elevations, making them priority areas for both research and conservation (Janzen, 1967; Rahbek et al., 2019). Moreover, tropical montane ecosystems are highly vulnerable to climate change, particularly for ectothermic taxa with narrow thermal niches (Janzen, 1967; Colwell et al., 2008). Understanding variation in species richness and community structure along these gradients is therefore essential for uncovering the processes that drive uneven biodiversity distribution and for anticipating biodiversity responses to environmental change.

In tropical ecosystems, moth diversity consistently declines from mid-elevations upwards (Beck et al., 2017), but species richness patterns in the lower parts of elevational gradients are more variable. Unimodal patterns with mid-elevation peaks predominate, as shown for geometrid moths in Costa Rica (Brehm et al., 2007), fruit-feeding moths on Mount Cameroon (Maicher et al., 2020a), sphingid moths in Sulawesi (Beck & Kitching, 2009), geometrid moths in Papua New Guinea (Toko et al., 2023), and large-bodied moths in Australia (Ashton et al., 2016). However, low-elevation peaks or low-elevation plateaus with a mid-elevation peak (terminology follows McCain & Grytnes, 2010) have also been reported, for example in most moth groups on Mount Cameroon (Maicher et al., 2020a) and in sphingid moths across several South-East Asian mountains (Beck & Kitching, 2009). In some cases, moth species richness declines linearly with elevation, as observed for various moth groups on Mount Kilimanjaro (Axmacher et al., 2004; Peters et al., 2016), while other studies report no consistent elevational pattern, such as for geometrid moths in Ecuador (e.g. Brehm et al., 2003). Micromoths (“Microleptidoptera”), a traditional polyphyletic group of small-bodied moths, remain particularly understudied, with the only elevational study from tropical mountains reporting a decline in pyraloid moths species richness with elevation in the Ecuadorian Andes (Fiedler et al., 2008).

Functional traits offer valuable insight into the mechanisms shaping biodiversity patterns along elevational gradients. Two traits particularly relevant in this context are species’ elevational range size and body size. The first is central to the elevational extension of Rapoport’s rule (Stevens, 1992), which predicts that species at higher elevations experience greater climatic variability and therefore evolve broader environmental tolerances and wider elevational ranges. Elevational range size is thus considered a proxy for climatic tolerance and dispersal capacity (Gaston, 2003). Empirical tests of Rapoport’s rule in tropical Lepidoptera have yielded mixed results: some studies report increasing range size with elevation (e.g. Brehm et al., 2007; Dewan & Acharya, 2024; Hnialum et al., 2025; Talukdar & Singh, 2025), whereas others find the opposite trend (e.g. Toko et al., 2023). Another widely discussed pattern is Bergmann’s cline (i.e. the elevational pattern of ectotherms’ body size consistent with predictions by Bergmann’s rule originally defined for endotherms; e.g. Blanckenhorn & Demont, 2004; Shelomi, 2012), which proposes that organisms attain larger body sizes in cooler environments, although the reverse pattern has often been proposed for ectotherms, including insects (e.g. Mousseau, 1997). In Lepidoptera, body size affects flight energetics and dispersal ability (Horne et al., 2017; Brehm et al., 2019), making it a key trait for assessing ecological strategies along elevational gradients. Although Bergmann’s cline has received relatively consistent support in tropical moths (e.g. Brehm et al., 2019; Mungee & Athreya, 2021), opposing patterns have also been reported (e.g. Brehm & Fiedler, 2004). Intraspecific analyses from Mount Cameroon similarly revealed both supporting (14 moth species) and opposing (5 moth species) patterns to Bergmann’s cline (Papandreou et al., 2023). Importantly, micromoths remain severely understudied in this context, with only a single elevational study documenting a linear decline in body size of Ecuadorian pyraloid moths with elevation (Scharnhorst & Fiedler, 2025).

Altogether, the available studies reveal no universal pattern of moth species richness along tropical elevational gradients, as observed trends vary across regions and moth lineages, probably reflecting interactions among ecological traits, regional history, and mountain-specific environmental filters. General trait-based patterns even less clear, as studies of Rapoport’s rule and Bergmann’s cline remain geographically patchy, strongly biased towards macromoths and the Neotropics or Indo-Malaya, and virtually absent for micromoths. This sparse and uneven coverage hampers robust comparative analyses of the underlying drivers shaping both species and trait distributions along tropical elevational gradients. Further research is urgently needed across underrepresented tropical regions and overlooked moth groups. In the Afrotropics, quantitative elevational data on moth communities (but not their traits) exist only for Mount Kilimanjaro and Mount Cameroon, and no study has yet examined elevational patterns in Alucitidae or Pterophoridae, the two focal micromoth families of this work, from any tropical region.

This study provides the first comprehensive analysis of micromoth diversity, community composition, and trait variation along an elevational gradient in the Afrotropics. Focusing on Alucitidae and Pterophoridae, two families with exceptional diversity and high endemism on Mount Cameroon (Ustjuzhanin et al., 2018, 2020, 2024, 2025), we address a major geographic and taxonomic gap in our understanding of elevational patterns in tropical insect diversity. Mount Cameroon’s continuous and largely intact forest gradient, spanning from sea level to the natural timberline, offers a unique opportunity to examine how species richness, community structure, and functional traits respond to steep climatic and environmental changes. Specifically, we (1) quantified patterns of species richness along the gradient, hypothesising a mid-elevation peak consistent with the most commonly observed patterns in other tropical Lepidoptera, while other patterns at low elevations can also be expected; (2) analysed changes in species composition with elevation, predicting marked turnover driven by strong environmental gradients; and (3) tested for elevational trends in species’ elevational range size and body size, hypothesising that both traits increase with elevation in accordance with Rapoport’s rule and Bergmann’s cline. By addressing these aims, our study provides the first functional-trait perspective on Afrotropical micromoth diversity.

## Methods

All analyses were performed in R 4.3.3 (R Core Team 2024).

### Study area

This study was conducted on Mount Cameroon, Cameroon (4.2167° N, 9.1700° E), the highest mountain in Western and Central Africa, reaching 4040 m a.s.l. The mountain is a recognised hotspot of biodiversity and endemism, particularly for Lepidoptera (e.g. Ustjuzhanin et al., 2018, 2020, 2024, 2025). The climate is strongly seasonal, with annual rainfall in the lowlands exceeds 12000 mm, including over 2000 mm per month during the wet season (June– September) and almost no precipitation in the dry season (mid-November–February; Maicher et al., 2018, 2020a).

### Micromoth diversity and trait datasets

We compiled records of Alucitidae and Pterophoridae from our earlier surveys of Mount Cameroon (Ustjuzhanin et al., 2018, 2020, 2024, 2025). Although these two micromoth families were not the primary targets of our previous ecological projects focused on moth and butterfly diversity on the mountain (see Maicher et al., 2020a,b), they were collected systematically, yielding a dataset suitable for diversity and trait analyses as described in Ustjuzhanin et al. (2018).

For this study, we included records of both focal groups from nine sampling sites on the southern and south-western slopes of Mount Cameroon, spanning seven elevations from littoral forest to the timberline (Table 1; further details in Maicher et al., 2020a,Maicher et al., 2020b). Moths were attracted to a light source emitting mainly in the white and UV spectrum (see Maicher 2020a for details) set at three plots per site. Sampling was carried out between 2014 and 2018 (exact dates for each site in Ustjuzhanin et al., 2018, Ustjuzhanin et al., 2020, Ustjuzhanin et al., 2024, Ustjuzhanin et al., 2025). Seven sites (BC-350, BB-30, CL-1500, DG-650, EC-1850, MS-2200, PC-1100) were sampled during three seasonal periods (transition from wet to dry seasons, full dry season, and transition from dry to wet season), while two sites (EF-1100, MF-1850) were sampled during the two transition periods only. At each site and season, moths were sampled over two nights per plot (see Table 1 for the number of sampling nights per site).

**Table 1.** Overview of sites on Mount Cameroon sampled for the diversity of Alucitidae and Pterophoridae moths, with each site’s elevation (m a.s.l.), geographic coordinates, abbreviation used in the text and figures, brief habitat characterisation, total number of light-trapping nights (across all plots and seasonal periods), and both observed (i.e. number of captured species) and estimated (Chao1 estimator ± SD) species richness.

| Sampling site | Elevation | Coordinates | Abbreviation | Habitat type | #nights | Observed species richness |  | Estimated species richness |  |
| --- | --- | --- | --- | --- | --- | --- | --- | --- | --- |
|  |  |  |  |  |  | Alucitidae | Pterophoridae | Alucitidae | Pterophoridae |
| Bimbia-Bonadikombo Forest | 30 m | 3.9818°N, 9.2625°E | BB-30 | Littoral forest, partly selectively logged | 18 | 2 | 1 | 3.0 ± 2.0 | 1.0 ± 0.1 |
| Bamboo Camp | 350 m | 4.0879°N, 9.0505°E | BC-350 | Mosaic of primary and secondary lowland forest | 18 | 15 | 4 | 17.5 ± 3.1 | 5.5 ± 2.5 |
| Drink Gari | 650 m | 4.1014°N, 9.0610°E | DG-650 | Primary lowland forest | 18 | 7 | 1 | 13.3 ± 7.0 | 1.0 ± 0.0 |
| Ekonjo Forest | 1100 m | 4.0881°N, 9.1168°E | EF-1100 | Primary submontane forest | 12 | 4 | 3 | 7.0 ± 4.3 | 3.5 ± 1.2 |
| PlanteCam Camp | 1100 m | 4.1175°N, 9.0709°E | PC-1100 | Upland forest locally disturbed by elephants | 18 | 23 | 10 | 41 ± 13.1 | 31 ± 17.4 |
| Crater Lake | 1500 m | 4.1443°N, 9.0717°E | CL-1500 | Submontane forest locally disturbed by elephants | 18 | 4 | 17 | 4.2 ± 0.1 | 21 ± 4.3 |
| Elephant Camp | 1850 m | 4.1170°N, 9.0729°E | EC-1850 | Montane forest locally disturbed by elephants | 18 | 4 | 9 | 4.0 ± 0.1 | 9.0 ± 0.1 |
| Mapanja Forest | 1850 m | 4.1157°N, 9.1315°E | MF-1850 | Primary montane forest | 12 | 5 | 12 | 11.2 ± 7.0 | 12.6 ± 1.1 |
| Mann's Spring | 2200 m | 4.1428°N, 9.1225°E | MS-2200 | Montane forest close to the timberline | 18 | 2 | 14 | 3.0 ± 2.0 | 16.5 ± 3.1 |

Due to the relatively low number of recorded individuals and species, we pooled all moths captured at each site across all plots, sampling nights, and seasons. Moreover, because of partly unbalanced sampling effort among sites, we did not use observed species richness (i.e. the number of captured species) in any analyses or direct comparisons. Instead, we calculated the Chao1 estimator, a bias-corrected, abundance-based estimate of species richness (Chiu et al., 2023), for each site, both for the combined dataset of both groups together and separately for each family. These estimates were obtained using the *estimateR* function in the *vegan* package (Oksanen et al., 2025) and are hereafter referred to as *estimated species richness*.

Each moth species was characterised by two traits. *Elevational range* was defined as the difference between the maximum and minimum capture elevations recorded in our dataset. *Wingspan*, as a widely accepted proxy for lepidopteran body size (Brehm et al., 2019; Mertens et al., 2021), was compiled from published sources (mainly Ustjuzhanin et al., 2018, 2020, 2024, 2025; De Prins & De Prins 2025) or measured directly from voucher specimens in the collections of P.U. and V.N.K. For species with known variability in wingspan, arithmetic means were used in all analyses. Trait values for all species are provided in Table S1. For each site, we then derived community-weighted means (CWMs) of elevational range and wingspan, calculated for both groups together and separately for each family.

### Statistical analyses

To visualise how species richness accumulated with sampling effort at each site, we constructed individual-based rarefaction curves using the *iNEXT* function with 1000 bootstrap replications in the *iNEXT* package (Hsieh et al., 2016). Curves were generated for both groups together and separately for each moth family, enabling visual evaluation of sampling completeness, particularly the tendency of curves to approach asymptotes, and qualitative comparison of species richness across sites while accounting for unequal numbers of individuals collected.

We then analysed the effect of elevation on estimated species richness and on CWMs of elevational range and wingspan. Each response variable was analysed with a separate linear model, applied to both groups together and each family individually, using *lm* function in the *stats* package (R Core Team, 2024). All response variables were visually checked for normality, resulting in square-root transformation of estimated species richness, whilst non-transformed CWMs of elevational range and wingspan were used in the models. For estimated species richness, we fitted both linear and quadratic (second-order polynomial) terms for elevation, whereas only a linear term was tested for CWMs of elevational range and wingspan.

To assess changes in community composition along the elevational gradient, we performed non-metric multidimensional scaling (NMDS) based on Bray–Curtis dissimilarity of species abundances at individual sampling sites. NMDS was conducted for both groups together and for each family seaparately, using the *vegan* package. To evaluate the effect of elevation on community composition, we used the *envfit* function with 999 permutations.

## Results

The analysed dataset comprised 471 micromoth specimens representing 64 species: 190 individuals of 31 Alucitidae species and 281 individuals of 33 Pterophoridae species (Fig. 1). Observed and estimated species richness of both groups at each site are presented in Table 1, and species abundances per site in Table S1. Fig. 1 also illustrates the contrasting elevational distributions of the two families. Most Alucitidae species were recorded at low to mid elevations (0–1100 m a.s.l.), with only a few species recorded at higher sites, whereas Pterophoridae species were more diverse and abundant from 1500 m a.s.l. upward.

**Figure 1.**
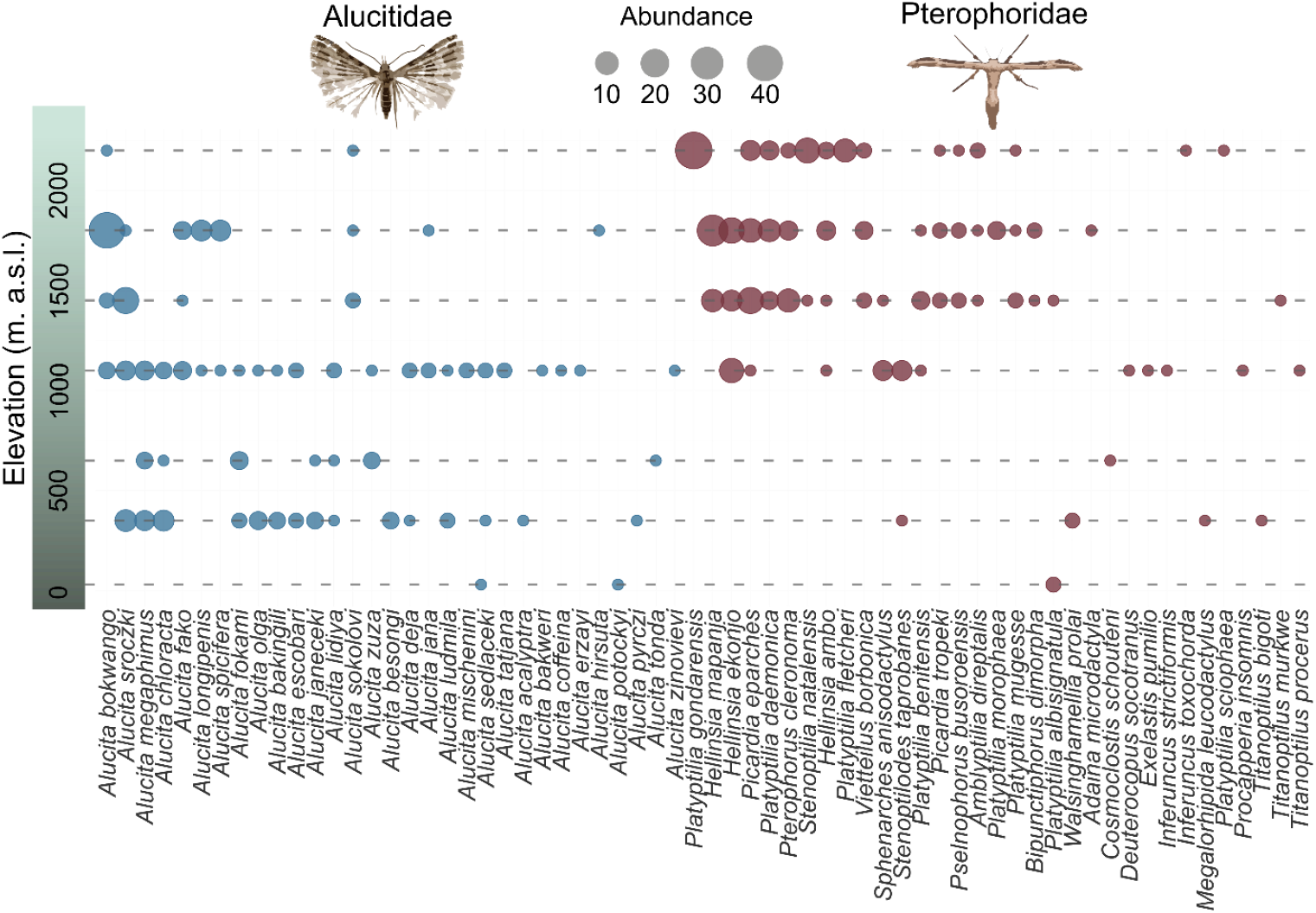
Elevational distribution and local abundance of the 64 micromoth species recorded on Mount Cameroon. Each column represents one species (Alucitidae = blue, Pterophoridae = maroon) ordered left to right by decreasing total abundance across all sites. Circles indicate each elevation where a species was recorded (y-axis), with circle size scaled to the number of individuals captured.

Rarefaction curves (Fig. 2) indicated that the mid-elevation plot PC-1100 consistently supported the highest extrapolated species richness for both groups together (Fig. 2a), and separately for Alucitidae (Fig. 2b) and Pterophoridae (Fig. 2c). Differences among the remaining sites were less pronounced: extrapolations for the adjacent high-elevation sites (CL-1500 and EC-1850) were closely clustered, while extrapolated species richness was notably lower at both the lowest site (BB-30) and the uppermost site (MS-2200). The curves for BB-30 and MS-2200 flattened early and showed narrow 95% confidence intervals, indicating near-complete sampling, whereas the curves for PC-1100 continued to rise and retain wide confidence bands, suggesting that additional species remain undetected at this elevation. Similarly, both observed and estimated species richness values (Table 1) consistently identified PC-1100 as the richest site for both Alucitidae and Pterophoridae, while BB-30 was the poorest for both families. The largest gap between observed and estimated species richness appeared at PC-1100, especially for Pterophoridae, again indicating incomplete sampling despite high observed diversity. In contrast, sites like EC-1850 and DG-650 showed only minimal differences between observed and estimated values, confirming the near-complete sampling.

**Figure 2.**
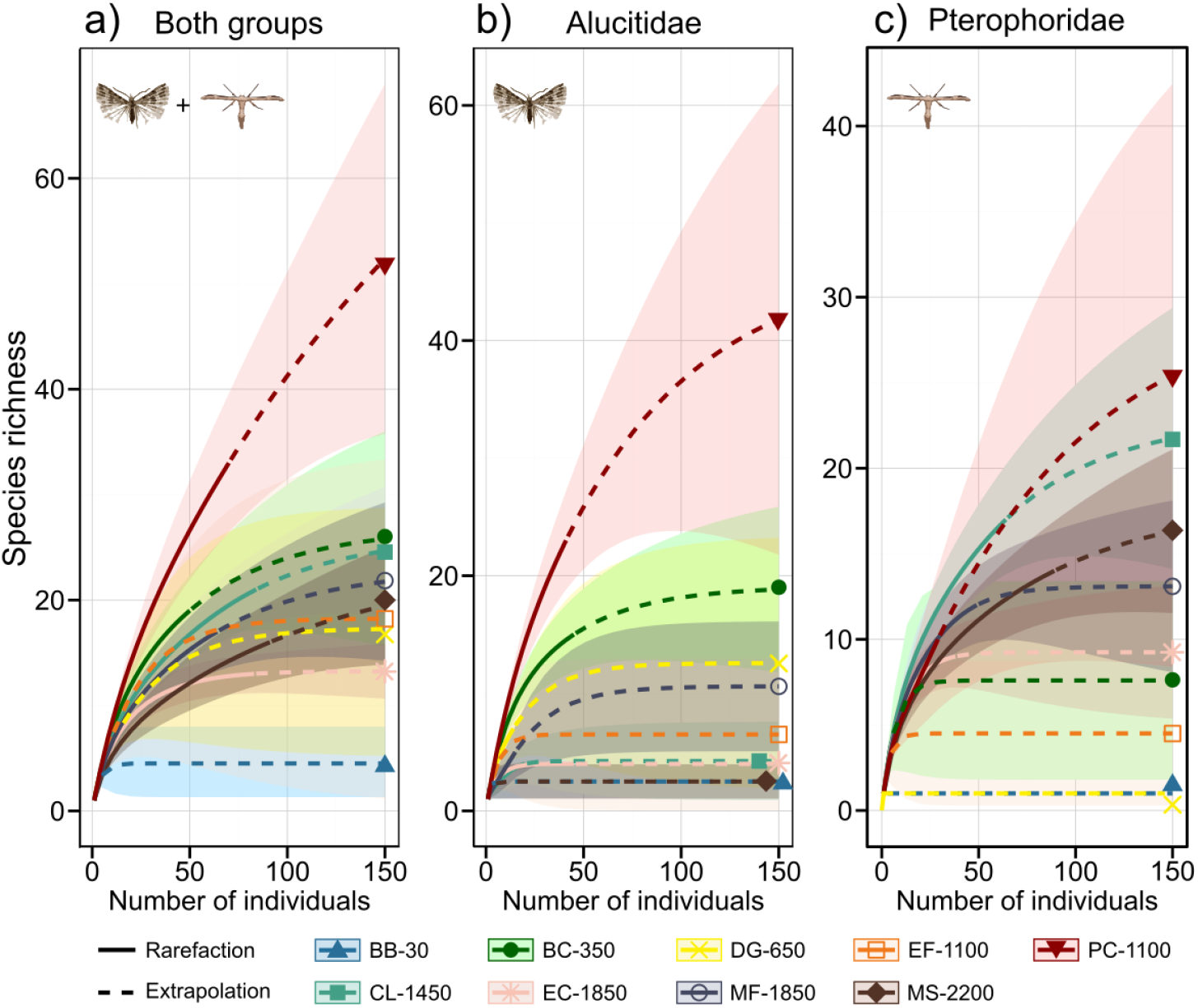
Individual-based rarefaction (solid lines) and extrapolation (dashed lines) curves with 95% confidence intervals for micromoth communities at individual sampling sites on Mount Cameroon: (a) both groups together, (b) Alucitidae, and (c) – Pterophoridae. Curves are colour-coded by site.

Linear models revealed contrasting elevational patterns in species richness and trait means (Fig. 3, Table 2). Estimated species richness showed a significant unimodal relationship with elevation for both groups together, peaking slightly above 1100 m a.s.l. (Fig. 3a). A similar hump-shaped pattern was found for Alucitidae, although the quadratic term was only marginally significant (Fig. 3a). In contrast, estimated species richness of Pterophoridae increased nearly linearly with elevation, lacking a mid-elevation peak (Fig. 3a). CWMs of elevational range size (Fig. 3b) and wingspan (Fig. 3c) increased significantly with elevation in all models (Table 2), consistent with Rapoport’s rule and Bergmann’s cline, respectively.

**Table 2.** Summary of linear model results testing the effects of elevation on estimated species richness (Chao1 estimator) and community-weighted means of elevational range size and wingspan in micromoths on Mount Cameroon, analysed for both moth groups together, and separately for Alucitidae and Pterophoridae. Models of estimated species richness included both linear and quadratic terms for elevation, while trait models included only linear terms. Model coefficients (Estimate), standard errors (SE), p-values (significant values in bold), and model fit (R^2^ and adjusted R^2^) are listed.

| Group | Term | Estimate | SE | p-value | $R^2$ | $R^2_{adj}$ |
| --- | --- | --- | --- | --- | --- | --- |
| *Estimated species richness* |  |  |  |  |  |  |
| Both groups | Elevation | 0.17 | 0.295 | 0.448 | 0.597 | 0.460 |
|  | Elevation<sup>2</sup> | -0.66 | 0.247 | **0.036** |  |  |
| Alucitidae | Elevation | -0.33 | 0.276 | 0.266 | 0.259 | 0.126 |
|  | Elevation<sup>2</sup> | -0.67 | 0.322 | 0.061 |  |  |
| Pterophoridae | Elevation | 0.82 | 0.352 | **0.045** | 0.534 | 0.380 |
|  | Elevation<sup>2</sup> | -0.31 | 0.403 | 0.468 |  |  |
| *Community-weighted mean elevational range* |  |  |  |  |  |  |
| Both groups | Elevation | 0.695 | 0.047 | **<0.001** | 0.968 | 0.964 |
| Alucitidae | Elevation | 0.649 | 0.040 | **<0.001** | 0.973 | 0.970 |
| Pterophoridae | Elevation | 0.721 | 0.092 | **<0.001** | 0.897 | 0.883 |
| *Community-weighted mean wingspan* |  |  |  |  |  |  |
| Both groups | Elevation | 0.0002 | 0.001 | **<0.001** | 0.857 | 0.836 |
| Alucitidae | Elevation | 0.0002 | 0.001 | **0.008** | 0.794 | 0.764 |
| Pterophoridae | Elevation | 0.0001 | 0.001 | **0.030** | 0.542 | 0.477 |

**Figure 3.**
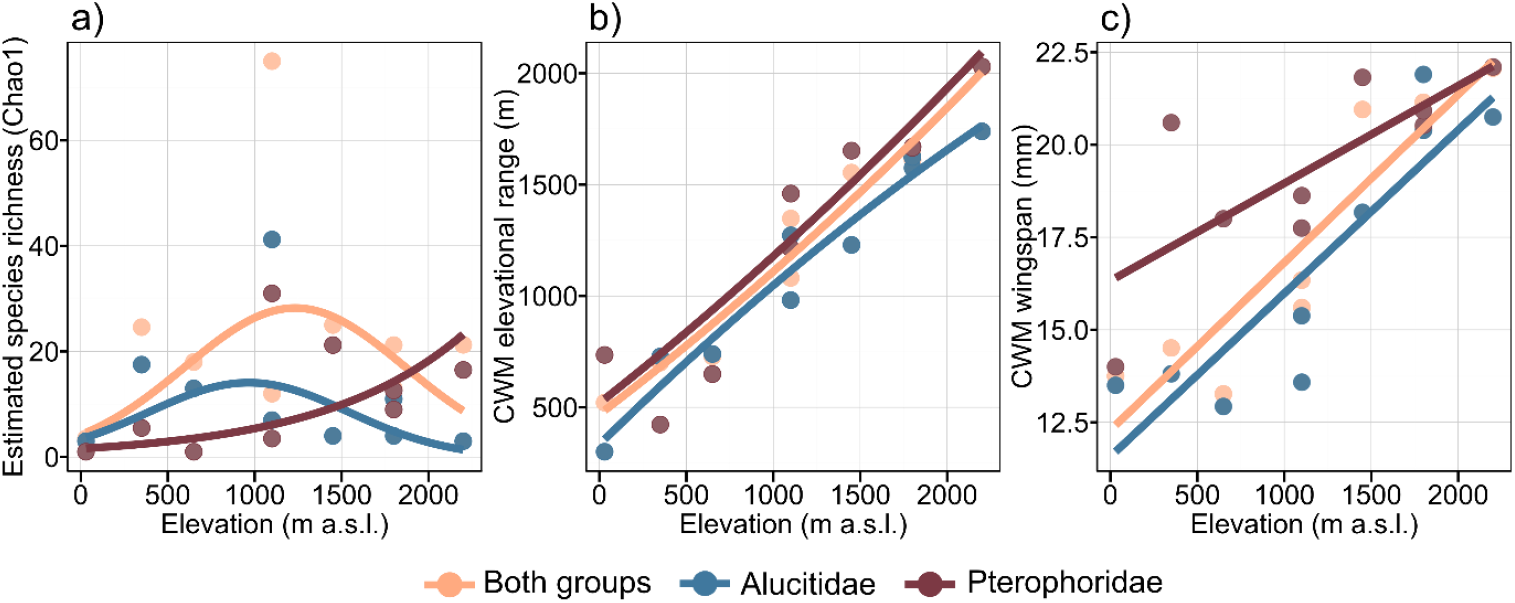
Elevational patterns for (a) estimated species richness, (b) community-weighted mean of elevational range size, and (c) community-weighted mean of wingspan in micromoth communities on Mount Cameroon. Points represent site-level values; coloured lines indicate significant linear models fitted for both groups together (orange), Alucitidae (blue), and Pterophoridae (red). Model statistics are provided in Table 2.

Species composition varied significantly with elevation for both groups together (*R*^*2*^ = 0.50, p < 0.05; Fig. 4a), and even more strongly when Alucitidae (*R*^*2*^ = 0.84, p < 0.01; Fig. 4b) and Pterophoridae (*R*^*2*^ = 0.77, p < 0.01; Fig. 4c) were analysed separately. For Alucitidae, sampling sites were clearly arranged along the first NMDS axis, reflecting a gradual turnover from lowland to montane communities, with the lowest site (BB-30) forming a distinct outlier (Fig. 4b). Pterophoridae showed a similarly strong elevational gradient, though primarily along the second NMDS axis, suggesting that additional environmental drivers may shape their community structure (Fig. 4c). When both families were analysed together, the elevational pattern was less distinct but still evident, with lowland sites clustering separately from mid- and high-elevation communities (Fig. 4a).

**Figure 4.**
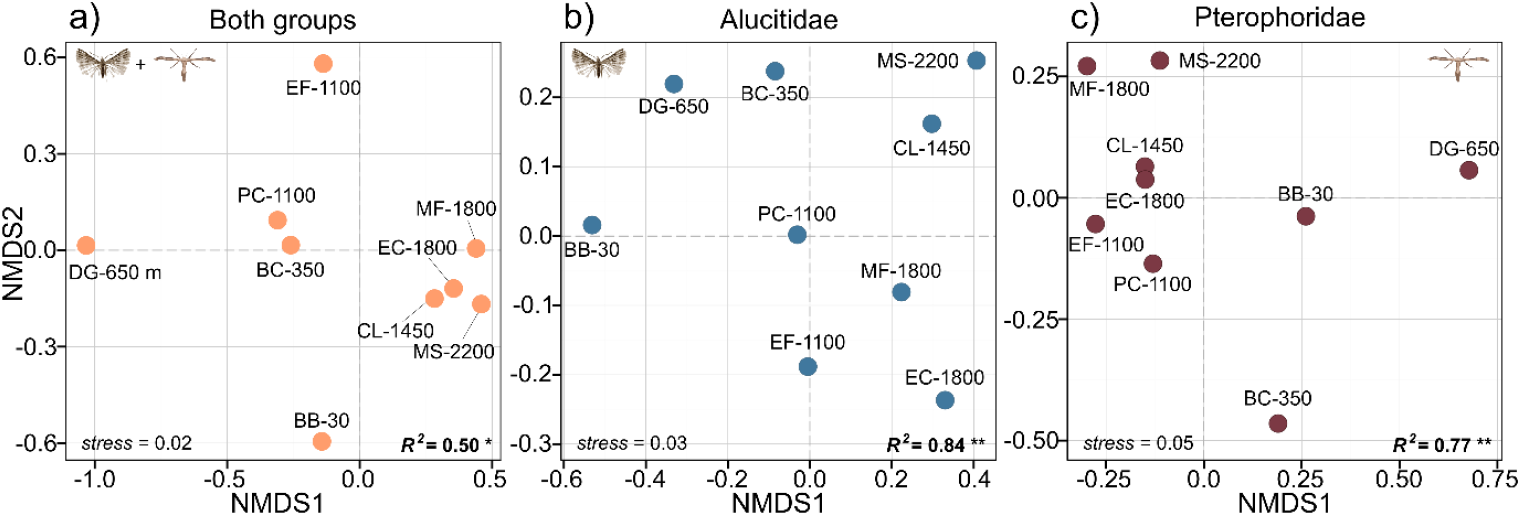
Non-metric multidimensional scaling (NMDS) ordinations of micromoth communities on Mount Cameroon: (a) both groups together, (b) Alucitidae, and (c) Pterophoridae. Points represent individual sampling sites (see Table 1 for the abbreviations). Stress values are shown in the lower left corner of each panel; R^2^ values from the envfit analyses for elevation are shown in the lower right. Significance levels: ***p < 0.001, **p < 0.01, *p < 0.05.

## Discussion

Our first comprehensive analysis of micromoth diversity along an Afrotropical elevational gradient, and one of the very few such studies from any tropical region, revealed contrasting elevational patterns between the two studied moth families. Alucitidae displayed a clear mid-elevation peak in species richness at 1100 m a.s.l., while Pterophoridae species richness increased with elevation. In both families, community composition shifted markedly along the gradient, reflecting strong environmental turnover. Furthermore, community-weighted elevational range size and wingspan both increased consistently with elevation, providing support for Rapoport’s rule and Bergmann’s cline in tropical micromoths.

Our findings on species richness patterns only partly align with previous elevational studies of tropical moths. The mid-elevation diversity peak of Alucitidae corresponds with the commonly reported unimodal patterns in many tropical moth groups (e.g. Brehm et al., 2007; Beck & Kitching, 2009; Ashton et al., 2016; Toko et al., 2023), including fruit-feeding moths on Mount Cameroon (Maicher et al., 2020a). However, most other moth groups on Mount Cameroon did not exhibit a similarly distinct mid-elevation peak: several peaked at lower elevations, while others showed broad low-elevation plateaus in species richness (*sensu* McCain & Grytnes, 2010) followed by declines in the uppermost forest zones (Maicher et al., 2020a; Mertens et al., 2021). In contrast, Pterophoridae species richness increased steadily with elevation, a pattern not documented in any other moth group on Mount Cameroon (Maicher et al., 2020a), nor elsewhere in tropical ecosystems, and opposite to the monotonic declines reported for several Afrotropical moth groups on Mount Kilimanjaro (Axmacher et al., 2004; Peters et al., 2016) and for pyraloid moths in the Ecuadorian Andes, which remains the only published elevational study on tropical micromoths (Fiedler et al., 2008). Nevertheless, that study reported increasing diversity with elevation for several pyraloid subfamilies (Scopariinae, Odontiinae, Galleriinae, Phycitinae; Fiedler et al., 2008), and similar elevational patterns have been observed in other non-tropical insect groups, such as ground beetles in Arizona (Uhey et al., 2022), and in tropical vertebrates, such as rodents in Tanzania (Stanley & Hutterer, 2007).

The observed elevational patterns of both micromoth groups are likely shaped by strong environmental filtering, as indicated by the pronounced elevational turnover of community composition. Both families consistently exhibited low species richness at the lowest elevations, with diversity increasing towards the mid- and/or upper slopes. Rarefaction curves and richness estimators confirmed that sampling was comparably effective across all elevations, with no indication that the lowland diversity results from undersampling. This shared pattern may thus reflect multiple environmental constraints in the lowlands. Littoral forests at the base of Mount Cameroon harboured the lowest lepidopteran species richness across previously studied groups (Maicher et al., 2020a), despite their high importance for species-rich communities of birds and trees (Ferenc et al., 2018). Their distinctive micromoth communities confirmed in our ordination analyses confirmed their high conservation importance for unique communities of other lepidopteran groups. The extremely high rainfall in the foothills (exceeding 12000 mm annually; Maicher et al., 2020a) may directly or indirectly limit survival of these small and fragile moths, whose minute, plume-like wings are especially vulnerable to sustained downpours. Additionally, strong predation pressure in lowland forests may disproportionately affect adults, while larvae of both families, often concealed borers in plant tissues (Gielis, 2003), may be less exposed. Predation pressure in tropical lowland forests is intense and declines with elevation (Roslin et al., 2017). Although insectivorous bird species richness peaks at mid-elevations on Mount Cameroon (Sedláček et al., 2023), this is likely offset by higher ant predation in the lowlands, a general pattern in tropical forests (Sam et al., 2015).

The mid-elevation peak in Alucitidae species richness may reflect a mid-domain effect or a local maximum in environmental heterogeneity, both commonly proposed drivers of such elevational diversity patterns (e.g. Colwell et al., 2016; Dolson & Kharouba, 2024; Camacho et al., 2025). Although the mid-domain effect is a widely recognised potential driver (e.g. Letten et al., 2013; Colwell et al., 2016), its role in shaping moth diversity on Mount Cameroon has not yet been explicitly tested. In contrast, forest structural heterogeneity is known to increase markedly with elevation on Mount Cameroon (Gaona et al., 2025), potentially creating a greater diversity of niches. Yet the Alucitidae diversity peak around 1100 m a.s.l., below the elevational zone of maximum heterogeneity (Gaona et al., 2025), suggests that other interacting drivers also contribute. Notably, the elephant-disturbed forest at mid elevation (PC-1100) hosted far more Alucitidae species than the adjacent undisturbed site (EF-1100), indicating that increased canopy openness from elephant activity (Maicher et al., 2020b) may enhance habitat suitability. However, this effect alone is insufficient to explain the elevational diversity peak, given that species richness was similar between disturbed (EC-1850) and undisturbed (MF-1850) sites at 1850 m a.s.l. Finally, climatic conditions likely play a role, as 1100 m a.s.l. coincides with a transitional climatic zone characterised by relatively moderate temperatures and rainfall (Maicher et al., 2020a).

The highly unusual increase in Pterophoridae diversity along elevation is difficult to compare with other studies. A comparable upslope pattern reported in some Ecuadorian pyraloid subfamilies was attributed to larval feeding on mosses and ferns more abundant in montane forests (Fiedler et al., 2008), but such host plants are rarely used by Pterophoridae moths (Matthews & Lott, 2005). Although host plants of Afrotropical Pterophoridae remain largely unknown, species from other regions often feed on various herbs (Matthews & Lott, 2005), which may be more abundant under the open canopy of montane forests. Comparable elevational diversity patterns in East African rodents have been linked to long-term climatic stability of Afromontane ecosystems and associated niche specialisation (Stanley & Hutterer, 2007), but such historical explanations are unlikely on the geologically young and volcanically active Mount Cameroon (Marzoli et al., 2000). Likewise, high-elevation diversity peaks reported for empidid flies in Thailand (Chatelain et al., 2018) and ground beetles in Arizona (Uhey et al., 2022) have been associated with regional precipitation regimes that differ substantially from those on Mount Cameroon (Maicher et al., 2020a). Furthermore, in contrast to Alucitidae, whose community composition was primarily structured by elevation, Pterophoridae communities showed elevation-related turnover along the second ordination axis, suggesting a stronger influence of other, unidentified environmental drivers or stochastic processes. Differences in evolutionary history may also contribute, given the markedly higher proportion of endemic species in Alucitidae (Ustjuzhanin et al., 2018, 2020, 2024) than in Pterophoridae (Ustjuzhanin et al., 2025), although the lack phylogenetic data and potential sampling biases across the Afrotropics limit any firm conclusions. Ultimately, without basic biological knowledge on Afrotropical Alucitidae and Pterophoridae, such as host-plant associations, habitat affinities, life history traits, and phylogenetic relationships, many established explanations for elevational diversity patterns in Lepidoptera (Beck et al., 2017) remain largely speculative for these groups.

Our study provides rare evidence supporting two widely discussed macroecological patterns in tropical micromoths. The confirmation of the elevation-extended Rapoport’s rule (Stevens, 1992) aligns with most previous studies on tropical butterflies and moths (e.g. Brehm et al., 2007; Dewan & Acharya, 2024; Hnialum et al., 2025; Talukdar & Singh, 2025), and likely reflects broader climatic tolerances or enhanced dispersal capabilities of high-elevation species (Gaston, 2003). Similarly, the observed support for Bergmann’s cline confirms the prevailing elevational trend in Lepidoptera body size (Brehm et al., 2019; Mungee & Athreya, 2021), despite ongoing debate about its applicability to ectotherms (e.g. Mousseau, 1997). Extending previous intraspecific evidence from Mount Cameroon, where most species exhibited increasing wingspan with elevation (Papandreou et al., 2023), we demonstrated a consistent community-level pattern in the two micromoth families. These findings, however, contrast with the declining body size trend reported for Ecuadorian pyraloid moths, the only comparable study on tropical micromoths (Scharnhorst & Fiedler, 2025 Potential mechanisms for the observed size increase include balancing of energy supply and demand under cooler conditions (Horne et al., 2017), or improved flights efficiency due to larger wings in lower-density air at higher elevations (Brehm et al., 2019). Nevertheless, without detailed experimental or physiological data, these hypotheses remain speculative and the mechanisms unresolved.

Our results on elevational patterns of micromoth diversity highlight several important implications for biodiversity conservation along the elevational gradient of Mount Cameroon. A substantial proportion of the regionally unique Alucitidae diversity (Ustjuzhanin et al., 2018, 2020, 2024) is concentrated at mid elevations, shaped by complex combinations of climatic and environmental drivers whose interactions remain poorly understood. In contrast, species-rich Pterophoridae communities, with a lower proportion of endemics compared to Alucitidae (Ustjuzhanin et al., 2025), dominate at higher elevations, although the specific ecological mechanisms behind their upslope diversity remain unclear. While species richness was comparatively low in the littoral forests at the lowest elevations, their distinct micromoth communities underline the high ecological value of these unique ecosystems, which are largely excluded from Mount Cameroon National Park and are increasingly threatened by logging and agricultural expansion (Ferenc et al., 2018). The observed elevational trait patterns suggest differential adaptive capacities among species. The presence of endemic species in both micromoth groups at high elevations further underscores their heightened extinction risk due to climate-driven habitat contraction (e.g. Morueta-Holme, 2015), while the lowlands may be particularly vulnerable to biological attrition caused by upslope range shifts and local extinctions under climate warming (e.g. Colwell et al., 2008). Together, these findings stress the need for targeted conservation strategies across the entire elevational gradient on Mount Cameroon. Simultaneously, we call for further research of the ecological and evolutionary drivers of diversity in neglected insect groups, particularly in tropical mountains where baseline data remain critically scarce. Such efforts, especially when supported by repeated long-term surveys, will be essential for predicting and mitigating the impacts of climate change on tropical biodiversity.

## Acknowledgements

We are grateful to Francis E. Luma, Jan E.J. Mertens, Jennifer T. Kimbeng, Ishmeal N. Kobe, Congo S. Kulu, and several other assistants for their help in the field, and to Eric B. Fokam and the staff of Mount Cameroon National Park for their support. We used ChatGPT (models o3 and 4o; OpenAI) for English proofreading. This study was conducted under authorisations issued by the Ministries of Forestry and Wildlife and of Research and Innovations of the Republic of Cameroon. The research was funded by the Czech Science Foundation.

## Data Availability Statement

All data used in this study are included in its supplementary materials.

