## Supplementary material for "Elevational patterns in two groups of micromoths (Lepidoptera: Pterophoridae, Alucitidae) in tropical forests of Mount Cameroon": Table S1

**Table S1.** Species occurrences and mean wingspan for Alucitidae and Pterophoridae micromoths across nine sampling sites on Mount Cameroon. Site codes (with elevation in m a.s.l.) are: BB-30 (Bimbia-Bonadikombo Forest, 30 m), BC-350 (Bamboo Camp, 350 m), DG-650 (Drink Gari, 650 m), EF-1100 (Ekonjo Forest, 1100 m), PC-1100 (PlanteCam Camp, 1100 m), CL-1500 (Crater Lake, 1500 m), EC-1850 (Elephant Camp, 1850 m), MF-1850 (Mapanja Forest, 1850 m), and MS-2200 (Mann's Spring, 2200 m). Numbers in the site columns indicate the individuals recorded.

| Family | Species | BB-30 | BC-350 | DG-650 | EF-1100 | PC-1100 | CL-1500 | EC-1850 | MF-1850 | MS-2200 | Wingspan (mm) |
|---|---|---|---|---|---|---|---|---|---|---|---|
| Pterophoridae | *Adaina microdactyla* | | | | | | | | 1 | | 15.0 |
| Pterophoridae | *Amblyptilia direptalis* | | | | | | 1 | 1 | | 2 | 13.0 |
| Pterophoridae | *Bipunctiphorus dimorpha* | | | | | | 1 | | 2 | | 16.0 |
| Pterophoridae | *Cosmoclostis schouteni* | | 1 | | | | | | | | 18.0 |
| Pterophoridae | *Deuterocopus socotranus* | | | | | 1 | | | | | 10.5 |
| Pterophoridae | *Exelastis pumilio* | | | | | 1 | | | | | 15.0 |
| Pterophoridae | *Hellinsia ambo* | | | | 1 | | 1 | 3 | 2 | 2 | 19.5 |
| Pterophoridae | *Hellinsia ekonjo* | | | | 2 | 11 | 7 | 9 | 4 | | 17.0 |
| Pterophoridae | *Hellinsia mapanja* | | | | | | 9 | 8 | 17 | | 18.0 |
| Pterophoridae | *Inferuncus strictiformis* | | | | | 1 | | | | | 18.0 |
| Pterophoridae | *Inferuncus toxochorda* | | | | | | | | | 1 | 12.0 |
| Pterophoridae | *Megalorhipida leucodactylus* | | 1 | | | | | | | | 19.0 |
| Pterophoridae | *Picardia eparches* | | | | | 1 | 15 | 3 | 8 | 6 | 24.5 |
| Pterophoridae | *Picardia tropeki* | | | | | | 2 | 2 | | 1 | 17.0 |

| | | | | | | | | | | | |
|---|---|---|---|---|---|---|---|---|---|---|---|
| Pterophoridae | *Platyptilia albisignatula* | 2 | | | | | 1 | | | | 14.0 |
| Pterophoridae | *Platyptilia benitensis* | | | | | 1 | 4 | | 1 | | 17.0 |
| Pterophoridae | *Platyptilia daemonica* | | | | | | 3 | 7 | 2 | 5 | 28.5 |
| Pterophoridae | *Platyptilia fletcheri* | | | | | | | | | 10 | 19.0 |
| Pterophoridae | *Platyptilia gondarensis* | | | | | | | | | 41 | 22.0 |
| Pterophoridae | *Platyptilia morophaea* | | | | | | | | 4 | | 25.5 |
| Pterophoridae | *Platyptilia mugesse* | | | | | | 2 | | 1 | 1 | 37.0 |
| Pterophoridae | *Platyptilia sciophaea* | | | | | | | | | 1 | 19.5 |
| Pterophoridae | *Procapperia insomnis* | | | | | 1 | | | | | 18.0 |
| Pterophoridae | *Pselnophorus busoroensis* | | | | | 2 | | 2 | | 1 | 17.0 |
| Pterophoridae | *Pterophorus cleronoma* | | | | | | 10 | 3 | 2 | | 23.5 |
| Pterophoridae | *Sphenarches anisodactylus* | | | | 1 | 5 | 1 | | | | 21.0 |
| Pterophoridae | *Stenoptilia natalensis* | | | | | | 1 | | | 13 | 23.5 |
| Pterophoridae | *Stenoptilodes taprobanes* | | 1 | | | 6 | | | | | 17.0 |
| Pterophoridae | *Titanoptilus bigoti* | | 1 | | | | | | | | 31.0 |
| Pterophoridae | *Titanoptilus murkwe* | | | | | | 1 | | | | 36.0 |
| Pterophoridae | *Titanoptilus procerus* | | | | | 1 | | | | | NA |
| Pterophoridae | *Vietteilus borbonica* | | | | | | 2 | | 4 | 2 | 23.0 |
| Pterophoridae | *Walsinghamellia prolai* | | 2 | | | | | | | | 20.0 |
| Alucitidae | *Alucita acalyptra* | | 1 | | | | | | | | 13.5 |
| Alucitidae | *Alucita bakingili* | | 3 | | | 1 | | | | | 11.0 |
| Alucitidae | *Alucita bakweri* | | | | | 1 | | | | | 18.0 |
| Alucitidae | *Alucita besongi* | | 3 | | | | | | | | 9.0 |
| Alucitidae | *Alucita bokwango* | | | | | 3 | 2 | 27 | 11 | | 24.0 |
| Alucitidae | *Alucita chloracta* | | 7 | 1 | | 3 | | | | | 14.5 |
| Alucitidae | *Alucita coffeina* | | | | | 1 | | | | | 20.0 |
| Alucitidae | *Alucita deja* | | 1 | | | 2 | | | | | 14.0 |
| Alucitidae | *Alucita erzayi* | | | | | 1 | | | | | 14.0 |
| Alucitidae | *Alucita escobari* | | 2 | | | 2 | | | | | 15.0 |
| Alucitidae | *Alucita fako* | | | | 3 | 3 | 1 | 3 | | | 10.5 |
| Alucitidae | *Alucita fokami* | | 2 | 4 | | 1 | | | | | 12.5 |
| Alucitidae | *Alucita hirsuta* | | | | | | | | 1 | | 14.0 |
| Alucitidae | *Alucita jana* | | | | 1 | 1 | | | 1 | | 15.0 |
| Alucitidae | *Alucita janeceki* | | 3 | 1 | | | | | | | 11.0 |
| Alucitidae | *Alucita lidiya* | | 1 | 1 | | 1 | | | | | 14.5 |
| Alucitidae | *Alucita longipenis* | | | | | 1 | | 7 | | | 20.5 |
| Alucitidae | *Alucita ludmila* | | 2 | | | 1 | | | | | 17.0 |
| Alucitidae | *Alucita megaphimus* | | 6 | 3 | | 1 | | | | | 13.0 |
| Alucitidae | *Alucita mischenini* | 1 | | | | 2 | | | | | 13.0 |
| Alucitidae | *Alucita olga* | | 4 | | | 1 | | | | | 10.5 |
| Alucitidae | *Alucita potockyi* | 1 | | | | | | | | | 14.0 |
| Alucitidae | *Alucita pyrczi* | | 1 | | | | | | | | 18.0 |
| Alucitidae | *Alucita sedlaceki* | | 1 | | 1 | 1 | | | | | 14.0 |
| Alucitidae | *Alucita sokolovi* | | | | | | 2 | | 1 | 1 | 17.5 |
| Alucitidae | *Alucita spicifera* | | | | | 1 | | 7 | | | 12.0 |
| Alucitidae | *Alucita sroczki* | | 8 | | | 5 | 15 | | 1 | | 18.0 |
| Alucitidae | *Alucita tatjana* | | | | 1 | 1 | | | | | 21.0 |
| Alucitidae | *Alucita tonda* | | | 1 | | | | | | | 16.0 |
| Alucitidae | *Alucita zinovievi* | | | | | 1 | | | | | 18.0 |
| Alucitidae | *Alucita zuza* | | | 3 | | 1 | | | | | 12.0 |
